# Cross-attention PHV: Prediction of human and virus protein-protein interactions using cross-attention–based neural networks

**DOI:** 10.1101/2022.07.03.498630

**Authors:** Sho Tsukiyama, Hiroyuki Kurata

## Abstract

Viral infections represent a major health concern worldwide. The alarming rate at which SARS-CoV-2 spreads, for example, led to a worldwide pandemic. Viruses incorporate genetic material into the host genome to hijack host cell functions such as the cell cycle and apoptosis. In these viral processes, protein-protein interactions (PPIs) play critical roles. Therefore, the identification of PPIs between humans and viruses is crucial for understanding the infection mechanism and host immune responses to viral infections and for discovering effective drugs. Experimental methods such as yeast two-hybrid assays and mass spectrometry are widely used to identify human-virus PPIs, but these experimental methods are time-consuming, expensive, and laborious. To overcome this problem, we developed a novel computational predictor, named cross-attention PHV, by implementing two key technologies of the cross-attention mechanism and a one-dimensional convolutional neural network (1D-CNN). The cross-attention mechanisms were very effective in enhancing prediction and generalization abilities. Application of 1D-CNN to the word2vec-generated feature matrices reduced computational costs, thus extending the allowable length of protein sequences to 9000 amino acid residues. Cross-attention PHV outperformed existing state-of-the-art models using a benchmark dataset and accurately predicted PPIs for unknown viruses. Cross-attention PHV also predicted human–SARS-CoV-2 PPIs with area under the curve values >0.95.

## Introduction

Viral infections represent a major health concern worldwide. The alarming rate at which severe acute respiratory syndrome coronavirus 2 (SARS-CoV-2) spreads, for example, led to a worldwide pandemic. According to the World Health Organization, more than 280 million people have been infected with SARS-CoV-2, and 5 million people had died by December 2021 [1].

Viruses enter host cells by interacting with receptors on the plasma membrane or by inducing endocytosis or membrane fusion [2–4]. To create an environment that promotes clone proliferation, viruses then incorporate its genetic material into the host genome to hijack host cell functions such as the cell cycle and apoptosis [5–7]. In these viral processes, protein-protein interactions (PPIs) play critical roles. Therefore, the identification of human and virus PPIs is crucial for understanding the infection mechanism and host immune responses and for discovering effective drugs. Experimental methods such as yeast-to-hybrid assays and mass spectrometry are widely used to identify human-virus PPIs (HV-PPIs). However, these experimental methods are not suitable for measuring all protein pairs because they are time-consuming, expensive, and laborious. To complement these existing experimental methods, various computational approaches have been adapted. Many protein structure–based PPI predictors have been proposed [8–10], but these tools are limited to predicting proteins for which the structure is known. As amino acid sequences are abundant and accessible, sequence-based prediction methods using machine learning and deep learning approaches have received increased attention.

Eid et al. proposed a sequence-based HV-PPI prediction framework called Denovo, which exhibited biological soundness and robust predictions [11]. They employed two important methods in development of Denovo. First, they constructed test datasets that included PPIs of viruses that are taxonomically far from viruses involved in PPIs of the training dataset according to rank in a taxonomy tree. Second, they employed a negative sampling method called “dissimilarity-based negative sampling”. Generally, negative data are essential for training models, but databases including non-interacting protein pairs do not exist, as far as we know. Therefore, negative samples need to be generated artificially. Many studies have used randomly sampled protein pairs without any experimentally verified interactions as negative samples [12], but this resulted in the generation of numerous false-negative results [13]. To deal with this issue, a dissimilarity-based negative sampling method was used.

In the prediction of HV-PPIs using machine learning models, amino acid sequences were encoded based on physicochemical properties, domain profiles, and sequence composition. Zhou et al. and Alguwaizani et al. generated feature vectors using compositional information to build SVM models [14, 15]. Their methods, which were evaluated with Denovo’s datasets, predicted HV-PPIs with an accuracy (AC) of around 85%. Furthermore, to enable predictions involving unknown viruses, the models were evaluated with a test dataset that excluded the virus species employed in the training dataset.

It is difficult for classical machine learning methods to extract local sequence patterns because they do not directly encode amino acid sequence-order information. Deep learning–based models have overcome such problems, however. For example, Yang et al. embedded local features such as binding motifs into feature matrices and captured their patterns using a convolutional neural network (CNN) [16]. They applied two different transfer learning methods to improve the generalizability of the model. Liu-Wei et al. developed a CNN-based HV-PPI predictor called DeepViral [17] by using not only sequence data but also disease phenotypes such as signs and symptoms.

Recently, natural language processing (NLP) methods such as word2vec [18] and doc2vec [19] have been applied for various biological predictions, including DNA N6- methyladenine site prediction [20], bitter peptide prediction [21], therapeutic peptide prediction [22], and prediction of compound-protein interactions [23]. These methods employ unsupervised embedding techniques to vectorize documents and words. For the prediction of HV-PPIs, Yang et al. used doc2vec-based embedding to extract contextual information from amino acid sequences [24]. We developed the long short-term memory (LSTM)-PHV [25] by adopting word2vec to consider consecutive 4-mers of amino acid sequences as words and captured the represented contextual information using an LSTM-based neural network. We demonstrated that LSTM-PHV accurately predicted HV-PPIs, whereas LSTM-PHV exhibited high computational and memory costs due to the recurrent LSTM computations. These methods exhibited good PPI prediction performance, but there is still room for improvement in predicting PPIs of unknown virus species.

To overcome such problems, we developed a novel sequence-based HV-PPI prediction model named cross-attention PHV. Instead of recurrent computations, we used attention modules. In particular, we crossed the two attention modules to mutually consider the features of human and virus proteins with respect to each other. It should be noted that this is the first application of cross-attention to PPI prediction, to the best of our knowledge. Furthermore, we applied a one-dimensional-CNN (1D-CNN) approach to increase the calculation speed, which resulted in extending the allowable length of protein sequences for training to 9000 residues, whereas the allowable sequence length of previous approaches is limited to 1000 to 2000 residues. The proposed method outperformed state-of-the-art models on Denovo’s datasets and accurately predicted unknown viral HV-PPIs.

## Materials and methods

### Dataset construction

#### Denovo’s dataset

Denovo’s dataset was employed to construct the cross-attention PHV. It consists of a training dataset with 5020 positive and 4734 negative samples and an independent test dataset with 425 positive and 425 negative samples [11]. We removed from the training dataset HV-PPIs that involved proteins with non-standard amino acids. The resultant training dataset included 5016 positive samples and 4732 negative samples.

### Human–unknown virus PPI dataset

To verify whether our proposed model predicts PPIs involving unknown viruses, we constructed a human–unknown virus PPI (HuV-PPI) dataset consisting of datasets of three influenza viruses, H1N1, H3N2, and H5N1. The HuV-PPI datasets were composed of training datasets without any samples of influenza viruses and the independent test datasets that included only samples of the influenza viruses, as shown in Table 1. In each dataset, the same numbers of negative samples and positive samples were randomly selected. Each dataset was divided into training and validation datasets at a ratio of 4:1.

**Table 1.**
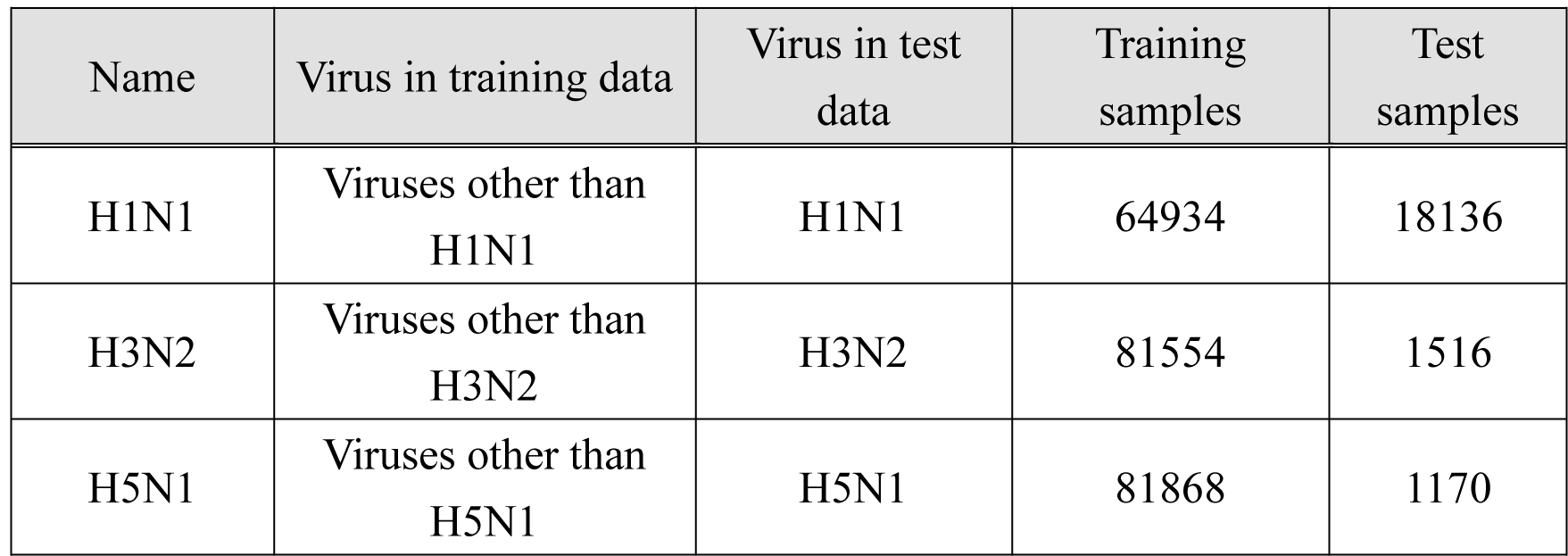
Statistical features of the HuV-PPI dataset.

| Name | Virus in training data | Virus in test data | Training samples | Test samples |
| --- | --- | --- | --- | --- |
| H1N1 | Viruses other than H1N1 | H1N1 | 64934 | 18136 |
| H3N2 | Viruses other than H3N2 | H3N2 | 81554 | 1516 |
| H5N1 | Viruses other than H5N1 | H5N1 | 81868 | 1170 |

Details regarding construction of the HuV-PPI datasets are described as follows. The PPIs and protein sequences were downloaded from the HVIDB [26] and UniProtKB (Swiss-Prot and TrEMBL) databases [27], respectively. We then removed the HV-PPIs involving proteins with a length of less than 30 or greater than 9000 residues and included non-standard amino acids. Negative samples were constructed using the dissimilarity-based negative sampling method in the same manner as our previous study [25]. This method uses a sequence similarity measure to explore protein pairs that are unlikely to interact. When virus protein A interacts with human protein B, we assumed that virus protein C, which exhibits less sequence similarity to virus protein A at an identity threshold of *T*, does not interact with human protein B. The pair of human protein B and virus protein C is thus a candidate negative sample. According to this approach, we compiled the negative PPI samples as follows. We combined human proteins registered in the UniProtKB/Swiss-Prot database with virus proteins included among the positive PPI samples. From the resulting human and virus protein pairs, we removed the pairs of the positive PPIs and further deleted the pairs of the human proteins and the virus proteins that exhibited higher similarity (*T* > 0.2) to the human protein-interacting virus proteins. We calculated the sequence similarities of all pairs of virus proteins included among the positive PPI samples using the Needleman-Wunsch algorithm with BLOSUM62 [28]. Note that human proteins with a length of less than 30 or greater than 9000 residues and proteins that included non-standard amino acids were removed.

### Human–SARS-CoV-2 PPI dataset construction

PPIs between human and SARS-CoV-2 were downloaded from the BioGRID database (COVID-19 Coronavirus Project 4.4.205) [29]. The sequence of each protein was retrieved from the UniProtKB database [27]. We removed PPIs involving proteins with a length of greater than 9000 residues or less than 30 residues and those that included non-standard amino acids. The remaining 14218 PPIs were used as positive samples. Negative samples were generated by applying the dissimilarity-based negative sampling method to human proteins retrieved from the UniProtKB/Swiss-Prot database [27]. The identity threshold was set to 0.2. The resultant dataset was named the human–SARS-CoV-2 PPI dataset. To build the balanced and imbalanced datasets, negative samples were randomly selected so that the ratios of positive to negative samples were 1:1 and 1:5, respectively. As shown in Table 2, the balanced and imbalanced datasets consisted of 28436 and 85308 samples, respectively. The resultant datasets were divided into training and test datasets at a ratio of 4:1.

**Table 2.**
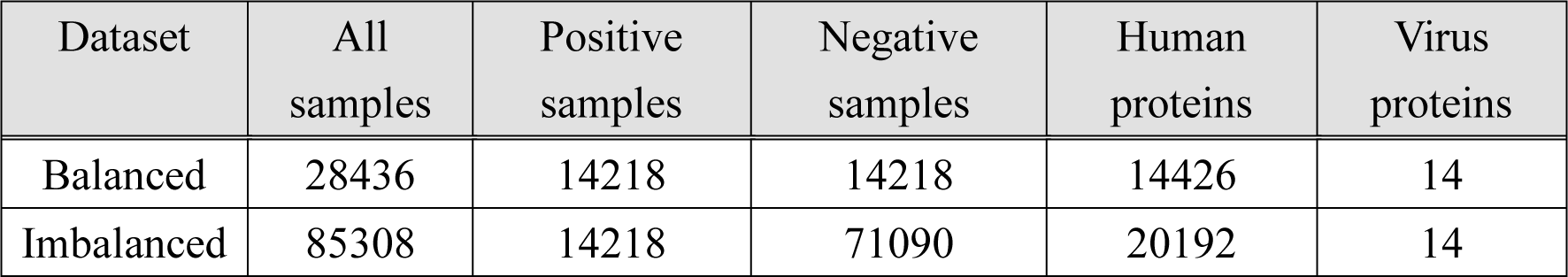
Statistical features of the SARS-CoV-2-PPI dataset.

### Feature encoding methods

The query sequences of human and virus proteins were encoded into feature matrices using word2vec, which generates distributed representations of words through a task that predicts a target word from its surrounding words (Continuous Bag-of-Words Model; CBOW) or predicts surrounding words from a target word (Continuous Skip-Gram Model; Skip-Gram). Although CBOW was employed in our previous study due to low computational cost [25], Skip-Gram was used in the present study because of its greater capacity to learn contextual information than CBOW [18]. Protein sequences were tokenized into consecutive k-mer amino acids (Fig. 1A).

**Fig. 1.**
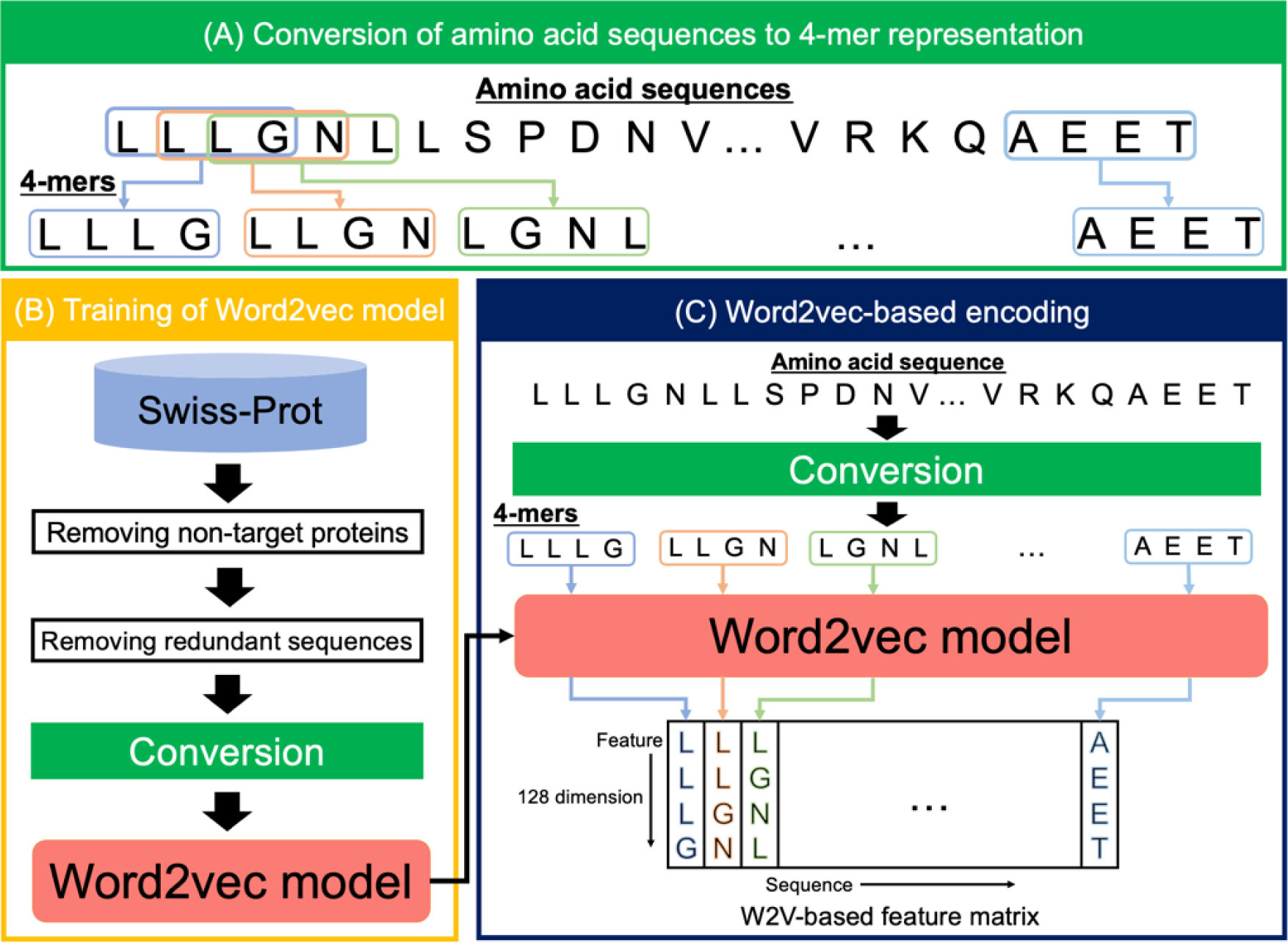
Workflow of word2vec-based encoding. (A) Amino acid sequences were converted into arrangements of consecutive 4-mers. (B) Amino acid sequences in the UniProtKB/Swiss-Prot database were converted as representations of 4-mers and used for training the word2vec model. (C) Each 4-mer in the amino acid sequence was converted into a feature vector using the trained word2vec model. The resultant feature vectors were concatenated into a feature matrix.

To train the word2vec model, we used the protein sequences in the UniProtKB/Swiss- Prot database [27] (Fig. 1B). Sequences with non-standard amino acids and those with a length greater than 9000 residues were excluded, and redundant sequences were then removed using CD-HIT with a threshold of 0.9 [30]. The remaining sequences were used to train the word2vec model. The maximum distance between a target k-mer and its surrounding k-mers (window size) and training iteration were set to 5 and 100, respectively.

Consecutive k-mer amino acids of protein sequences were transformed by the trained word2vec model into 128-dimensional feature vectors. These vectors were concatenated in the order of the sequences (Fig. 1C) to arrange the feature matrixes with a shape of (9000 − k+1) × 128, where zero-padding was applied so that the maximum sequence size was 9000. We constructed the word2vec model using Genism (version: 3.8.3) in the Python package (version: 3.8.0).

### Neural networks

#### 1D-CNN with max-pooling layer

To extract the features of local sequences such as binding motifs, 1-D convolutional layers were used. In these layers, input matrix *X* with *n* length and *s* channels was converted to feature matrix *C* with (*n* − *w*)⁄*t* + 1 length and *f* channels using sliding with shift width *t* and *f* filters with size *w*. The *i*-th element *C*_*i,k*_ of the matrix generated by the *k*-th filter *M*^k^ is given by:

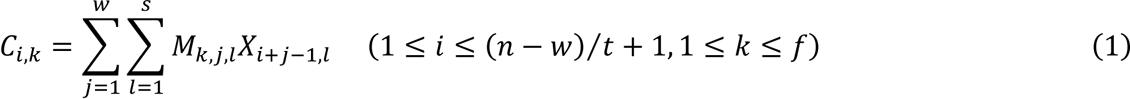

The pooling layer was placed at the position following the convolutional layers to suppress overfitting and increase generalization ability. The max-pooling layer samples the maximum values from the certain area of the input as follows:

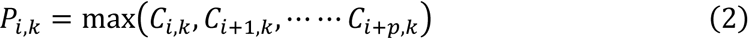

^where *p* represents the size of the pooling window. Zero-padding was applied to the^ input matrix so that the lengths of the input and output of the pooling layer were the same.

A global max-pooling layer generated a vector by sampling the maximum value from each channel of the output as follows:

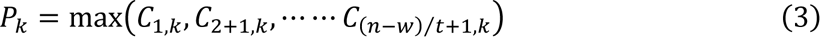

### Attention mechanism

Attention mechanism has been an important contributor to the remarkable advances that have occurred in neural network development, and it has been incorporated in recent neural network models such as BERT [31] and Transformer [32]. In the attention mechanism, output feature *Y*^*out*^ is generated by updating pre-updated feature *Y*^*pre*^ with information-giving feature *X*. For the update, three representations, known as Query, Key, and Value, are generated by applying three different learnable weights, *W*^*Q*^, *W*^*K*^, and *W*^*V*^, to these features, as follows:

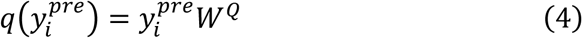

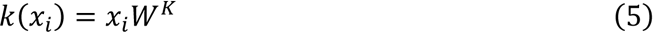

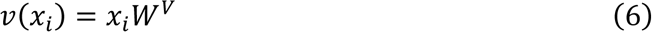

where *Y*^*pre*^_*i*_ and *x*_*i*_ indicate the *i*-th feature vectors of *Y*^*pre*^ and *X*, respectively, and *q*(·), *k*(·), and *v*(·) represent the transformation functions for calculating Query, Key, and Value, respectively. Next, attention weight α_*i,j*_, which determines the degree of influence of *x*_*j*_ on the calculation of updated vector *y*^*out*^_*i*_ of *y*^*pre*^_*i*_, is given by scaling the dot-product between Key and Query with dimension *d*_*key*_ of Key and by applying the masking and softmax functions to the scaled dot-product, as follows:

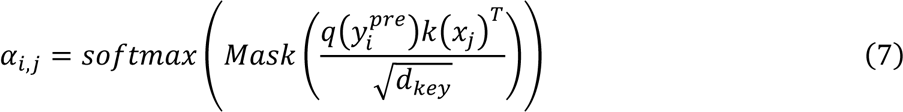

In the masking, the elements of padding position are set to minus infinity. Consequently, the effect of zero-padding can be neglected after applying the softmax function. To selectively extract information from the Value, depending on the relationship between

Key and Query, the weighted sum of Value is calculated with the attention weights, as follows:

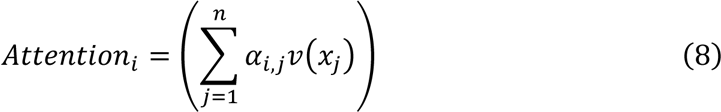

Next, the feature *Y*^*out*^ is updated as follows:

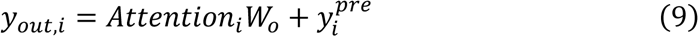

^where learnable weight *W*_*o*_ is applied to the weighted sum. In the multi-head attention^ layers, the weighted sums are calculated in parallel in each “head”, concatenated, and applied by *W*_*o*_.

### Cross-attention PHV

As shown in Figure 2, cross-attention PHV is composed of three sub-networks: (1) convolutional embedding modules, (2) a cross-attention network module, and (3) a feature integration network. The key technologies are to use 1D-CNN, which effectively extracts the features of long-length sequences of human and virus proteins and to develop a cross-attention module that extracts some feature interactions between human and virus proteins as the core of the learning method. Importantly, the attention modules represent human and virus proteins to capture global information regarding the amino acid sequences. We crossed the two attention modules to mutually consider the features of human and virus proteins.

**Fig. 2.**
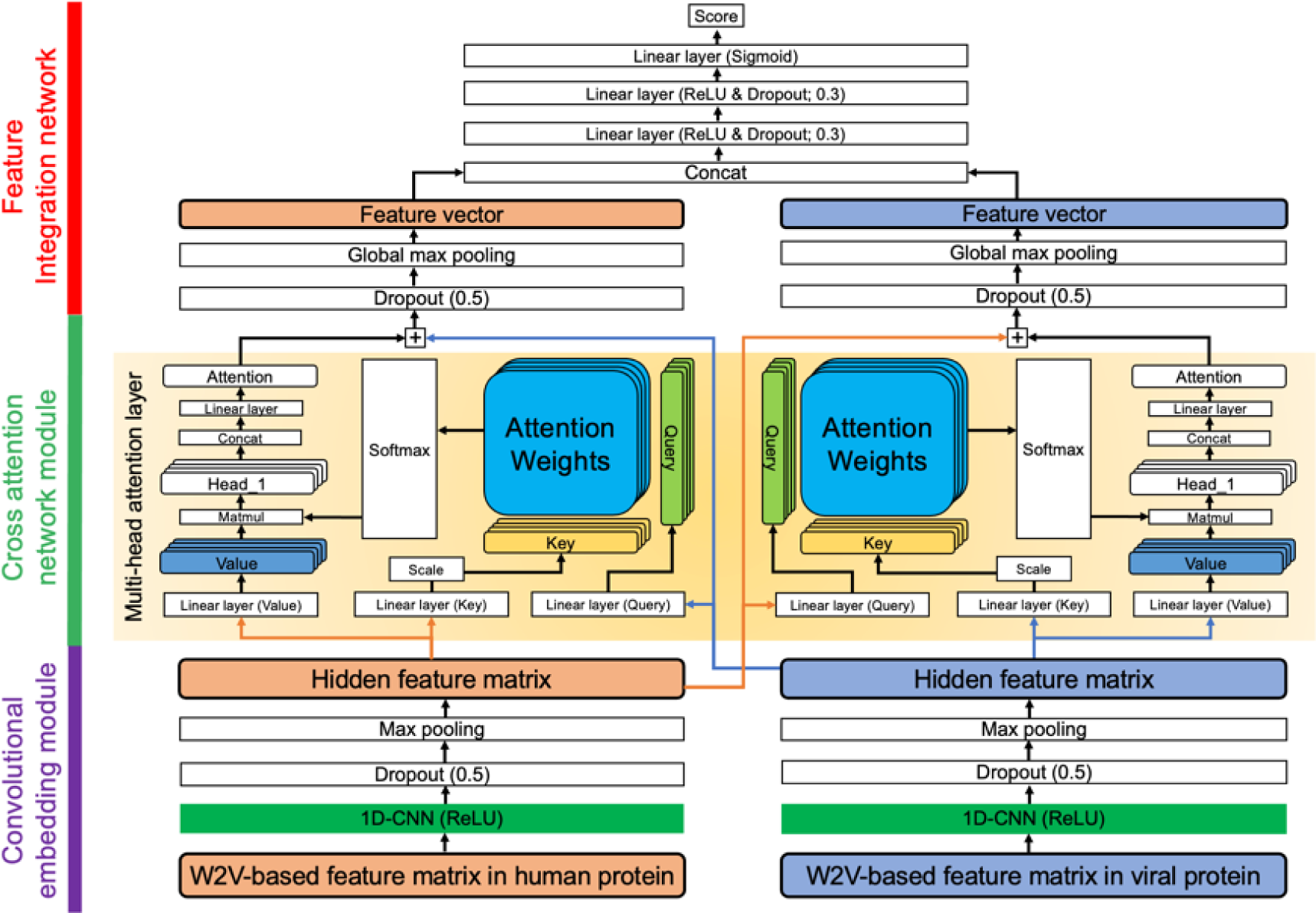
Structure of cross-attention PHV. Cross-attention PHV is composed of three sub-networks. The word2vec (W2V)-based feature matrices of humans and viruses were input into the convolutional embedding module. To extract interaction features between two protein sequences, multi-head attention layers were employed in the cross-attention module. Finally, the feature vectors generated by the global max-pooling layer were concatenated to compute a final score through three linear layers.

In the convolutional embedding modules, the word2vec-generated matrices of human and virus proteins were filtered by 1D-CNN layers with 128 filters and by max-pooling layers with a pooling window of 3. The size and shift width of the filters were set to 20 and 10, respectively. Based on a previous study [33], we inserted a dropout layer with a ratio of 0.5 between the 1D-CNN and the pooling layers. This transformation provides two advantages. One advantage is that it significantly decreases computational cost and memory usage for the next layers due to the reduced dimension of the feature matrix. Another advantage is that each vector of these feature matrices is generated from consecutive k-mer amino acids, which makes it possible to learn the dependencies among local patterns such as motifs.

In the cross-attention network module, the filtered matrices of human and virus proteins in the first sub-network were input into cross-attention modules consisting of two multi-head attention modules. We crossed the two attention modules to extract the features of human and virus proteins while referring to virus and human protein features, respectively. Specifically, one attention module applied the Query from the human feature matrix to the Keys from the virus feature matrix to calculate the attention weights, generating the attention of the Values of the virus feature matrix. This attention was used to extract virus protein features related to the Query from human protein features. In the same manner, the other attention module extracted the human protein features related to the virus protein features. The number of heads and the dimension of feature representations (Query, Key, Value) were set to 4 and 32, respectively.

In the feature integration network, the feature matrices processed by the cross-attention modules were transformed into feature vectors by global max-pooling layers having a dropout layer with a dropout ratio of 0.5. The feature vectors were then concatenated and transferred to the three fully connected layers to compute the final output. The hidden vectors from the first and second fully connected layers were dropped out at a ratio of 0.3. The vector dimensions from the first and second fully connected layers were set to 64 and 16, respectively. We constructed the whole neural network model using PyTorch in the Python package (version: 3.8.0).

### Training and testing

In the training scheme, loss was calculated using a binary cross-entropy function for each mini-batch of size 32. Optimization was executed using the Adam optimizer with a learning rate of 0.0001. To prevent over-learning, the training was stopped (early stopping) when the maximum area under the curve (AUC) was updated for 20 consecutive epochs.

### Measures

Six statistical measures were employed to evaluate the trained model: sensitivity (SN; recall), specificity (SP), accuracy (AC), Matthews correlation coefficient (MCC), F1- score (F1), and AUC. The formulas for calculating the measures other than AUC are given by:

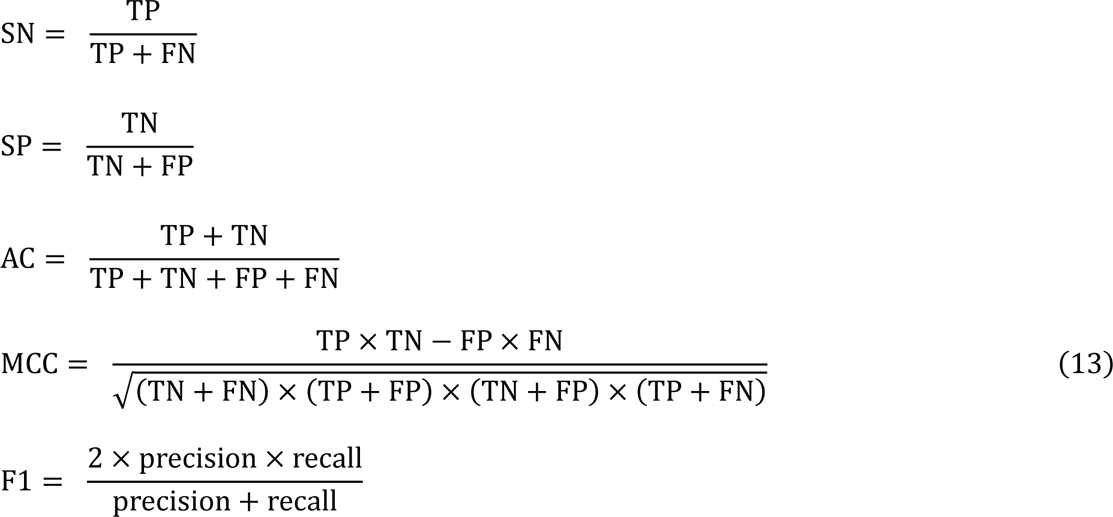

where TP, FP, TN, and FN indicate the numbers of true-positive, false-positive, true-negative, and false-negative samples, respectively. The threshold to determine whether proteins interact was set at 0.5. These measures were calculated using scikit-learn of the Python package [34].

### Visualization of features

To visualize feature vectors, we used t-distributed stochastic neighbor embedding (t-SNE) [35]. Compared with classical linear mapping methods such as principal component analysis and multiple discriminant analysis, t-SNE precisely projects both local and global structures of high-dimensional vectors into low-dimensional representations and is suitable for visualization of nonlinear data. The perplexity in t-SNE was set to 50.

## Results and Discussion

### Optimization of cross-attention PHV

We encoded the protein sequences into the feature matrices using the word2vec model and trained cross-attention PHV using Denovo’s training dataset. To achieve the best model, we optimized the k-mer value between 2 and 4 via 5-fold cross-validation, in which the training data were divided into 5 subsets, and then 4 subsets were used for training the model; the remaining subset was used for validation. Cross-attention PHV presented AUCs >0.97 on average, and the 4-mer model provided the highest values in terms of AC, MCC, AUC, and F1 (Fig. 3). Thus, we set the k-mer value to 4.

**Fig. 3.**
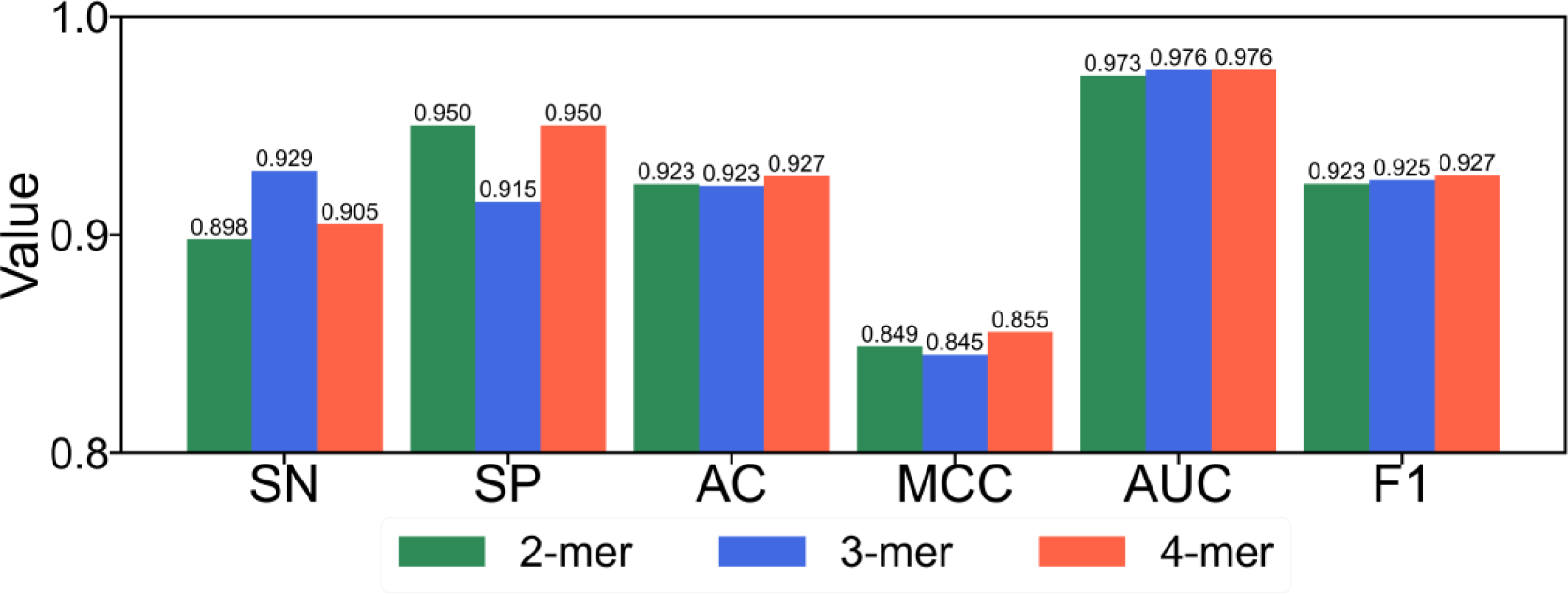
Prediction performance of word2vec-based cross-attention PHV with respect to k-mer value. The models were evaluated via 5-fold cross-validation on Denovo’s training dataset.

To demonstrate the superiority of word2vec, we used a binary encoding as the reference to construct the binary encoding–based cross-attention PHV. The binary encoding concatenated the one-hot vectors of each amino acid in the order of the protein sequence. The word2vec-based and binary encoding–based models were trained via 5- fold cross-validation on Denovo’s training dataset and evaluated using Denovo’s test dataset. To compare the two encoding methods, a two-sample *t*-test was applied to the AUC and AC values. As shown in Figure 4, the word2vec-based model (cross-attention PHV) provided significantly better performance than the binary encoding–based model (AUC; *p*-value <0.01, AC; *p*-value <0.05), suggesting that word2vec efficiently represents protein sequence contextual information.

**Fig. 4.**
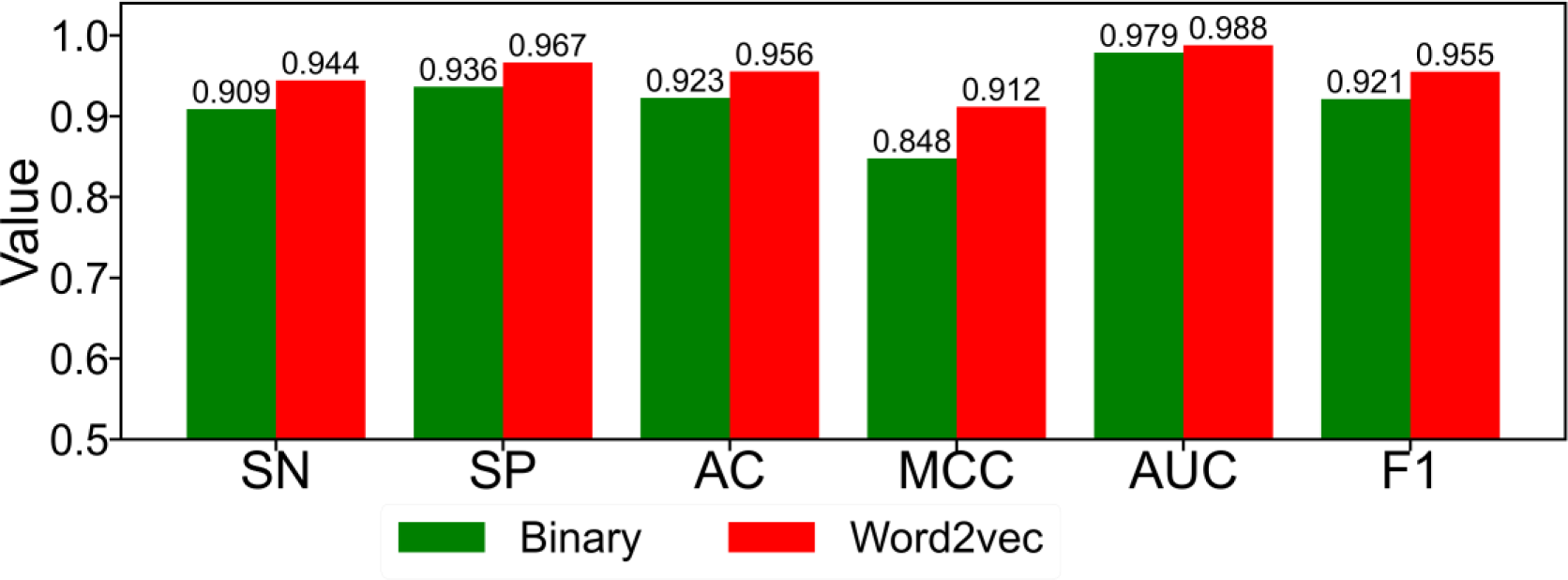
Comparison of performance between the word2vec-based and binary encodings in cross-attention PHV. Models trained via 5-fold cross-validation were evaluated with Denovo’s test dataset.

The cross-attention modules were expected to extract features of human and virus proteins by considering various relationships between local patterns in the two protein sequences. We compared cross-attention PHV with a self-attention–based neural network in which the features of human and virus proteins were input separately to multi-head attention modules without any interactions (Fig. S1). Both the cross-attention PHV and the self-attention–based neural network were trained via 5-fold cross-validation with Denovo’s training dataset. Cross-attention PHV outperformed the self-attention–based neural network (Fig. 5) on Denovo’s test dataset, suggesting that interactions between the features of human and virus proteins improve the prediction performance.

**Fig. 5.**
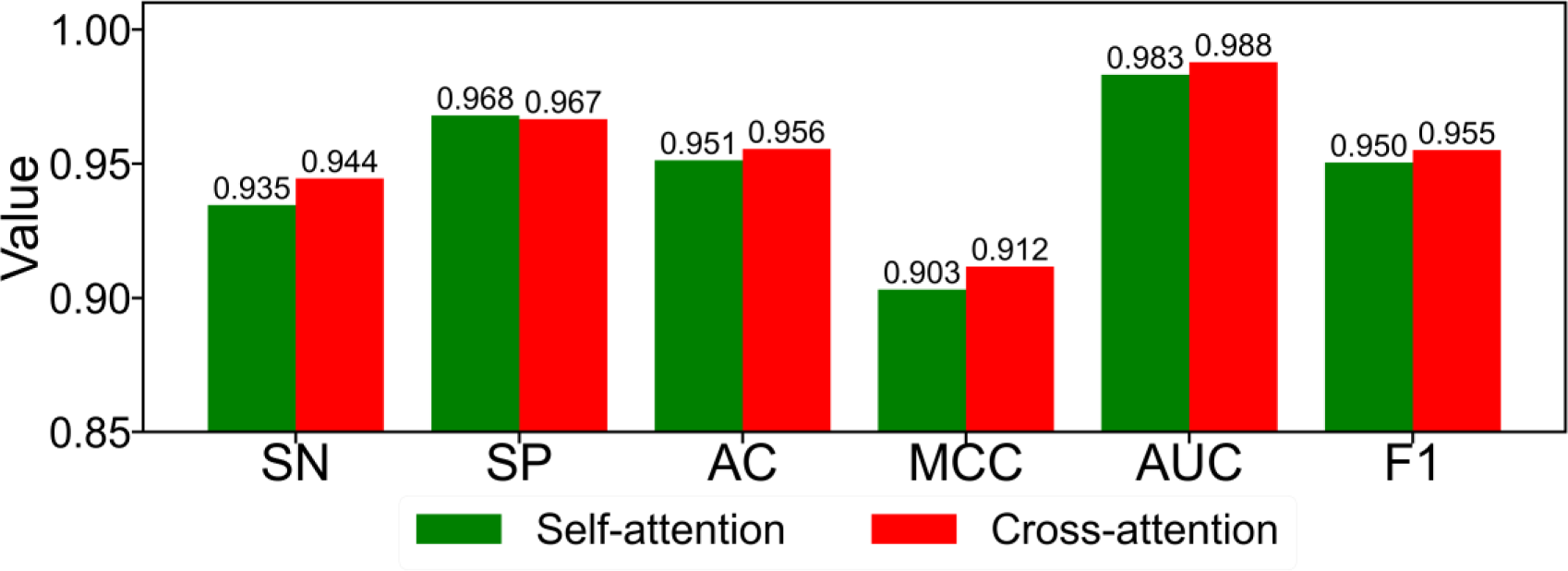
Comparison of performance between cross-attention–based and self-attention–based neural networks. Models trained via 5-fold cross-validation were evaluated with Denovo’s test dataset.

### Comparison of state-of-the-art methods

We compared the performance of cross-attention PHV with seven state-of-the-art methods, including Denovo [11], Zhou et al.’s SVM-based method [14], Alguwaizani et al.’s SVM-based method [15], Yang et al.’s random forest–based and Doc2vec-based method [24], DeepViral [17], and Yang et al.’s CNN-based method [16], using Denovo’s test dataset. As shown in Table 3, cross-attention PHV predicted the PPIs with an AC value >0.95 and outperformed the state-of-the-art models in five metrics, including SN, AC, AUC, MCC, and F1, demonstrating the superiority of cross-attention PHV. We attributed the high prediction performance of cross-attention PHV to the capturing of contextual, interrelated information between the two feature matrices of human and virus protein sequences.

**Table 3.**
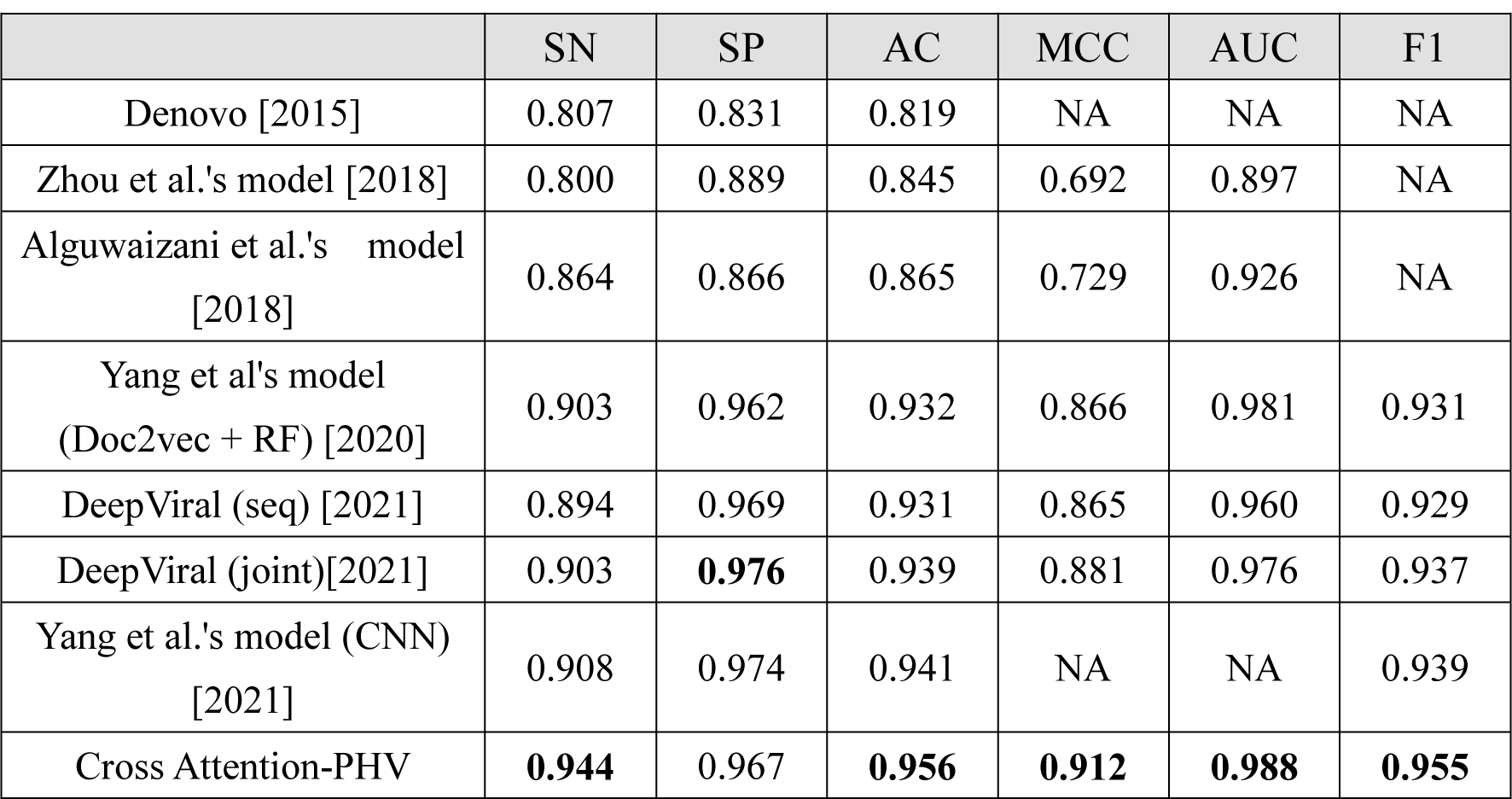
Comparison of the performance of cross-attention PHV with existing state-of- the-art models on Denovo’s test dataset. Data regarding the performance of existing models were obtained from the respective papers. Bold values indicate the highest value for each measurement.

### Performance in predicting PPIs between humans and unknown virus species

To evaluate the PPI prediction performance between humans and unknown viruses, we employed the HuV-PPI (H1N1, H3N2, and H5N1) datasets. It should be noted that the training dataset excluded the PPIs of the influenza species employed in the independent test dataset. Twenty percent of the training data were used for validation of early stopping. We compared cross-attention PHV with LSTM-PHV [25], which exhibited the best performance in the year 2021. In the training of LSTM-PHV, proteins with a sequence length greater than 1000 residues were removed in the same manner [25] because the method has a significant memory and time cost. As shown in Figure 6, cross-attention PHV exhibited AC and AUC values >0.91 and >0.96, respectively, on the independent datasets. Cross-attention PHV outperformed LSTM-PHV, demonstrating the high generalizability of cross-attention PHV. As the encoding method (word2vec) used in cross-attention PHV is the same as that of LSTM-PHV, the cross-attention–based network was found to be more predictive than the LSTM-based network. Furthermore, cross-attention PHV does not require the recursive calculations employed by LSTM-PHV, which greatly reduces computation time. This is also major advantage of cross-attention PHV, as it enables us to process long sequences for the training scheme.

**Fig. 6.**
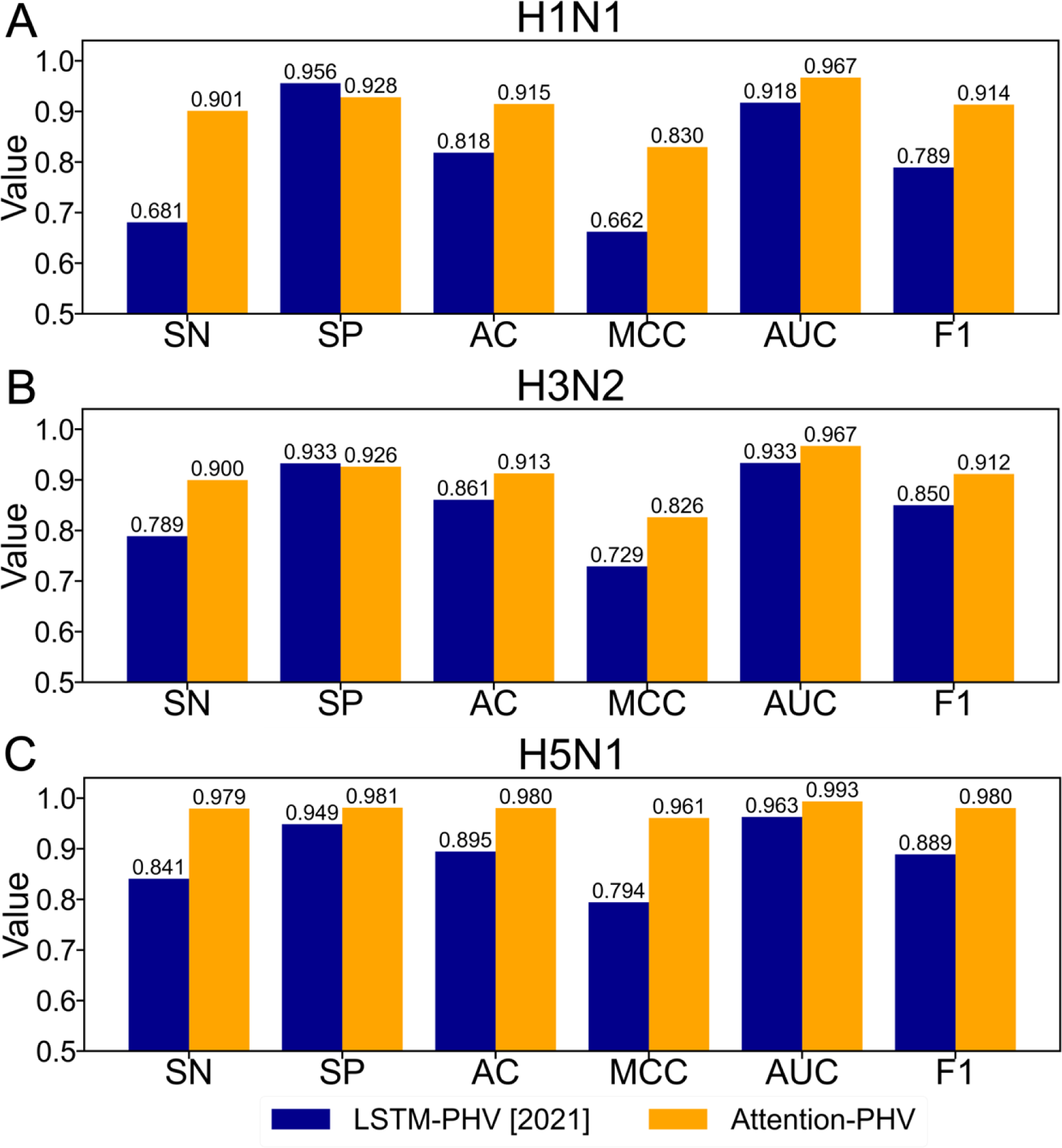
Comparison of the performance of cross-attention PHV and LSTM-PHV in predicting PPIs for unknown viruses. (A) Performance on the H1N1 dataset, which regards H1N1 as an unknown virus. (B) Performance on the H3N2 dataset, which regards H3N2 as an unknown virus. (C) Performance on the H5N1 dataset, which regards H5N1 as an unknown virus.

### Performance in predicting PPIs between humans and SARS-CoV-2

We also investigated whether cross-attention PHV can predict PPIs between humans and SARS-CoV-2 using the human–SARS-CoV-2 PPI dataset. The models were trained via 5-fold cross-validation with the training dataset and evaluated with its independent test dataset. All measures were averaged over the five models. We characterized cross-attention PHV in comparison with LSTM-PHV. In training the LSTM-PHV, we removed PPIs involving proteins with a sequence length greater than 1000 residues to minimize the computational cost. As shown in Figure 7, cross-attention PHV exhibited AUCs >0.95 with both the balanced and imbalanced datasets. Furthermore, cross-attention PHV exhibited better performance than LSTM-PHV for all measures except SN. In particular, when the model was trained on a limited number of negative samples in the balanced dataset, cross-attention PHV demonstrated higher generalization ability than LSTM-PHV.

**Fig. 7.**
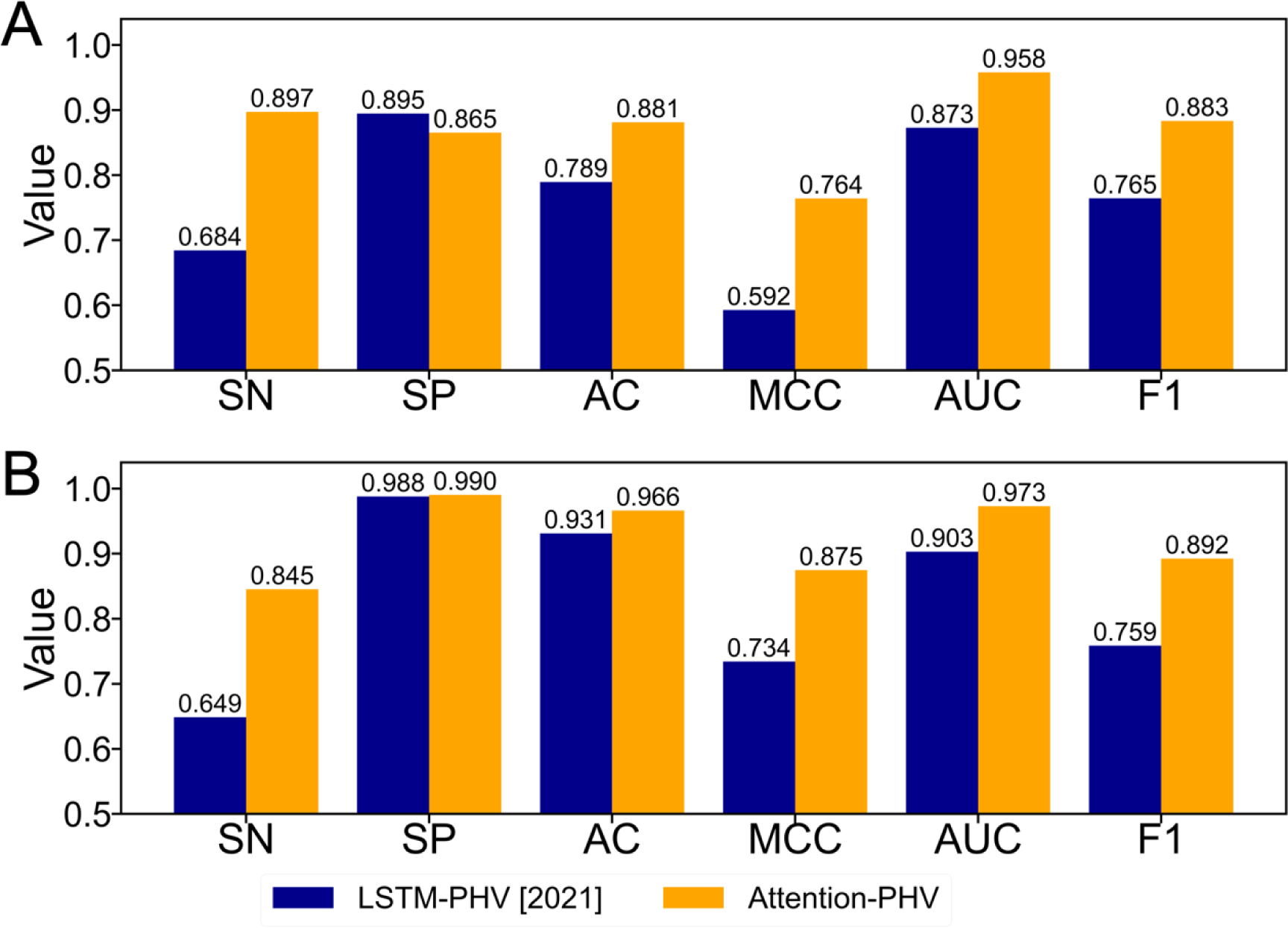
Comparison of the performance of cross-attention PHV with LSTM-PHV in predicting human–SARS-CoV-2 PPIs. (A) Performance on a balanced dataset (positive:negative = 1:1). (B) Performance on an imbalanced dataset (positive:negative = 1:5).

### Visualization and analysis of feature vectors and matrices

To investigate how each subnetwork of cross-attention PHV contributes to the prediction of PPIs between humans and unknown viruses, t-SNE was used to visualize the three features during testing of the HuV-PPI datasets: the word2vec-based feature matrices, hidden feature matrices, and feature vectors (Fig. 2). Before visualization, the word2vec- based feature matrices and hidden feature matrices were transformed into vectors by sampling the maximum values of each feature. The feature vectors for humans and viruses were then concatenated. As shown in Figure 8, for all the three datasets, the distributions of positive and negative PPI samples became clearly separated during testing, suggesting that the convolutional embedding and cross-attention modules extract important features responsible for the prediction. Furthermore, we visualized the concatenated human and virus feature vectors. Interestingly, the shapes of the feature vector distributions differed between humans and viruses (Fig. 9), reflecting the evolutionary or taxonomic differences between human and virus proteins.

**Fig. 8.**
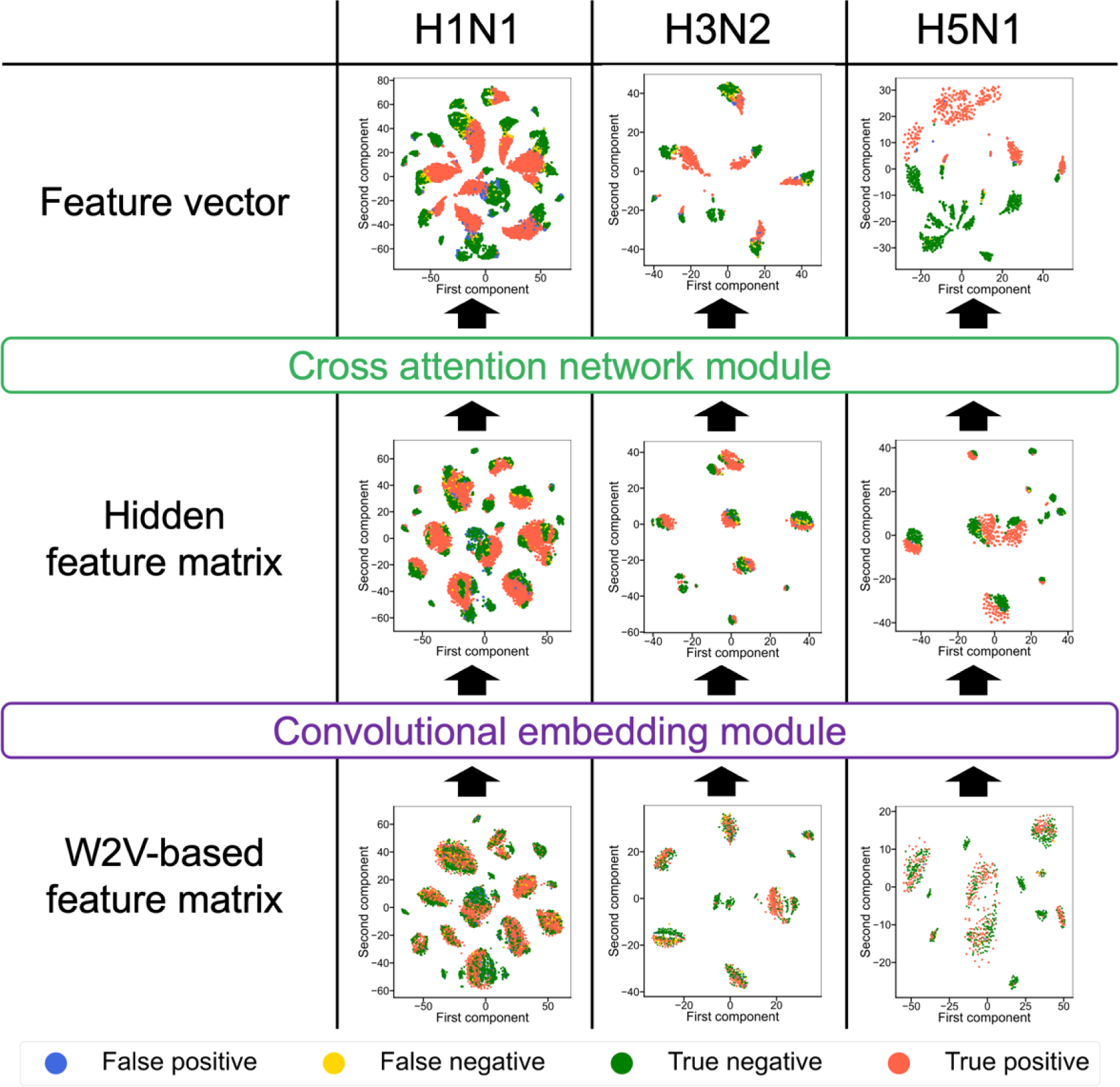
t-SNE–based visualization of features generated during prediction of PPIs using the HuV-PPI test datasets. The word2vec-based feature matrices, hidden feature matrices, and feature vectors were retrieved from the neural networks. The feature matrices were transformed into vectors by sampling the maximum values of each feature. The human and virus feature vectors were then concatenated. The t-SNE maps for the H1N1, H3N2, and H5N1 datasets are shown at the left, center, and right, respectively. Blue, yellow, green, and red marks indicate false-positive, false-negative, true-negative, and true-positive samples, respectively.

**Fig. 9.**
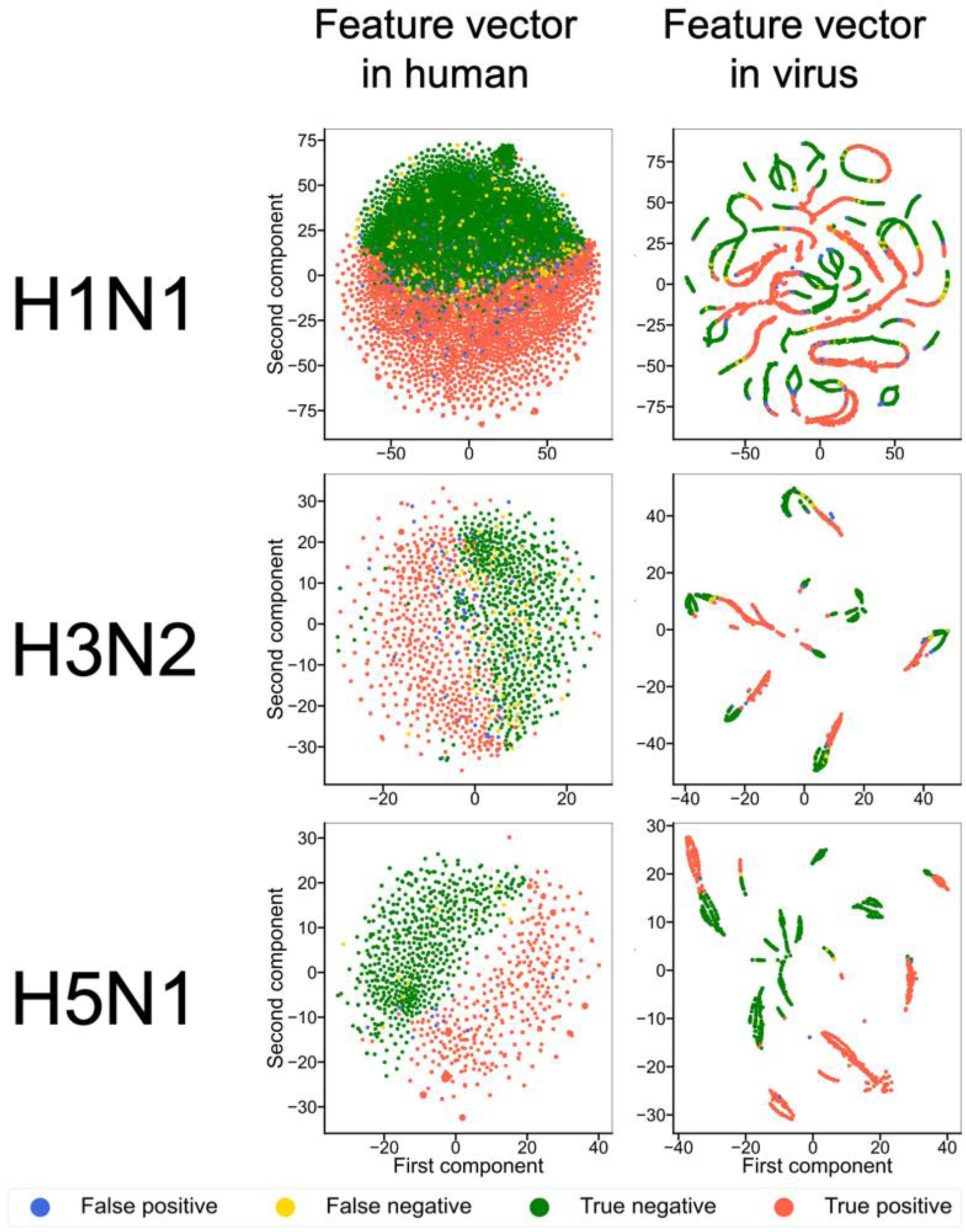
t-SNE–visualized map of the respective human and virus feature vectors on the HuV-PPI datasets.

### Limitation

Our proposed neural network was able to increase the allowable length of protein sequences to 9000 residues because a 1D-CNN was adopted to reduce the dimension of the sequences. However, such a feature extraction method makes it difficult for the attention weight to identify k-mer amino acid residues responsible for interactions. In future work, we hope to propose a methodology to identify amino acids important for interactions.

### Conclusions

To construct the cross-attention PHV predictor for PPIs between humans and viruses, we applied two key technologies, a cross-attention mechanism and a 1D-CNN. The cross-attention mechanism was very effective in achieving enhanced prediction and generalization to unknown virus species. Application of the 1D-CNN to word2vec- generated feature matrices extended the allowable length of protein sequences to 9000 residues for training scheme. We employed the word2vec model to embed the protein sequences and optimized the k-mer value of the word2vec model. Cross-attention PHV outperformed the state-of-the-art models for the five measures of SN, AC, MCC, AUC, and F1 using Denovo’s benchmark dataset. Furthermore, cross-attention PHV outperformed the best model of the year 2021 (LSTM-PHV) in predicting PPIs for unknown viruses. Finally, we demonstrated that cross-attention PHV captures the features responsible for virus infection–related proteins and distinguishes taxonomic and evolutionary differences between human and virus proteins.

## Acknowledgement

This work was supported by a Grant-in-Aid for Scientific Research (B) (22H03688) and partially supported by a Grant-in-Aid for JSPS Research Fellows (22J22706) from the Japan Society for the Promotion of Science (JSPS).

## Key points

- Cross-attention PHV implements two key technologies: (1) a cross-attention module to mutually consider the features of human and virus proteins; (2) a 1D-CNN to extend the allowable length of protein sequences up to 9000 residues.
- Cross-attention PHV outperforms state-of-the-art models in predicting PPIs between humans and viruses and accurately predicts PPIs for unknown influenza viruses and SARS-CoV-2.
- Cross-attention PHV extracts critical features responsible for PPI prediction and distinguishes taxonomic and evolutionary differences between human and virus proteins.

## Biographical note

**Sho Tsukiyama** is a PhD student in the Department of Creative Informatics in the Kyushu Institute of Technology, Japan. His main research interests include deep learning and computational biology.

**Hiroyuki Kurata** is a professor in the Department of Bioscience and Bioinformatics in the Kyushu Institute of Technology, Japan. His research interests primarily focus on systems biology, synthetics biology, functional genomics, machine learning, and their applications.

## Supplemental information

**Fig. S1.**
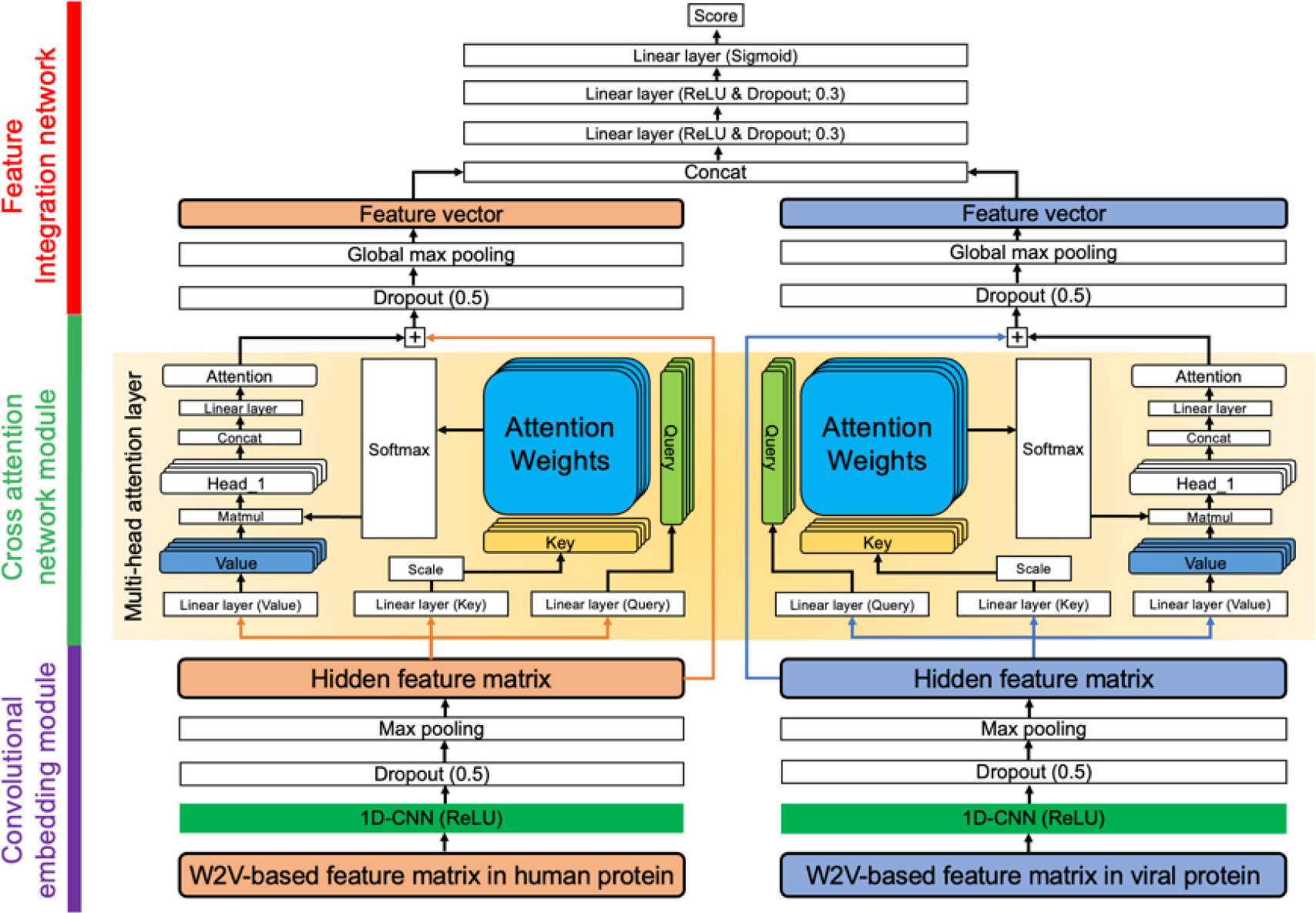
Structure of the self-attention–based neural network. The network was composed of three sub-networks. The word2vec (W2V)-based feature matrices of human and virus proteins were input into the convolutional embedding module. The output matrices of human and virus proteins were then input into the separate respective multi-head attention layers. This part differs from cross-attention PHV. Finally, the feature vectors generated by the global max-pooling layer were concatenated to compute a final score through three linear layers.

## Notes

### Competing Interest Statement

The authors have declared no competing interest.

## References

1. World Health Organization et al. Coronavirus disease (covid-19) situation dashboard. https://covid19.who.int/ (December 29 2021, date last accessed).

2. Burckhardt CJ, Greber UF. Virus movements on the plasma membrane support infection and transmission between cells, PLoS pathogens 2009;5:e1000621–e1000621.

3. Grove J, Marsh M. The cell biology of receptor-mediated virus entry, Journal of Cell Biology 2011;195:1071–1082.

4. Maginnis MS. Virus-Receptor Interactions: The Key to Cellular Invasion, J Mol Biol 2018;430:2590–2611.

5. Parnell G, McLean A, Booth D et al. Aberrant cell cycle and apoptotic changes characterise severe influenza A infection--a meta-analysis of genomic signatures in circulating leukocytes, PLoS One 2011;6:e17186–e17186.

6. Li FQ, Tam JP, Liu DX. Cell cycle arrest and apoptosis induced by the coronavirus infectious bronchitis virus in the absence of p53, Virology 2007;365:435–445.

7. Gioti K, Kottaridi C, Voyiatzaki C et al. Animal Coronaviruses Induced Apoptosis, Life 2021;11.

8. Hayashi T, Matsuzaki Y, Yanagisawa K et al. MEGADOCK-Web: an integrated database of high-throughput structure-based protein-protein interaction predictions, BMC Bioinformatics 2018;19:62.

9. Singh R, Park D, Xu J et al. Struct2Net: a web service to predict protein-protein interactions using a structure-based approach, Nucleic Acids Res 2010;38:W508–515.

10. Zhang QC, Petrey D, Deng L et al. Structure-based prediction of protein–protein interactions on a genome-wide scale, Nature 2012;490:556–560.

11. Eid FE, ElHefnawi M, Heath LS. DeNovo: virus-host sequence-based protein-protein interaction prediction, Bioinformatics 2016;32:1144–1150.

12. Barman RK, Saha S, Das S. Prediction of interactions between viral and host proteins using supervised machine learning methods, PLoS One 2014;9:e112034.

13. Dey L, Chakraborty S, Mukhopadhyay A. Machine learning techniques for sequence-based prediction of viral-host interactions between SARS-CoV-2 and human proteins, Biomed J 2020;43:438–450.

14. Zhou X, Park B, Choi D et al. A generalized approach to predicting protein-protein interactions between virus and host, BMC Genomics 2018;19:568.

15. Alguwaizani S, Park B, Zhou X et al. Predicting Interactions between Virus and Host Proteins Using Repeat Patterns and Composition of Amino Acids, Journal of Healthcare Engineering 2018;2018:1391265.

16. Yang X, Yang S, Lian X et al. Transfer learning via multi-scale convolutional neural layers for human–virus protein–protein interaction prediction, Bioinformatics 2021;37:4771–4778.

17. Liu-Wei W, Kafkas Ş, Chen J, et al. DeepViral: infectious disease phenotypes improve prediction of novel virus–host interactions, bioRxiv 2020:2020.2004.2022.055095.

18. Mikolov T, Chen K, Corrado G, et al. Efficient Estimation of Word Representations in Vector Space. 2013, arXiv:1301.3781.

19. Mikolov T, Sutskever I, Chen K, et al. Distributed Representations of Words and Phrases and their Compositionality. 2013, arXiv:1310.4546.

20. Tsukiyama S, Hasan MM, Deng H-W et al. BERT6mA: prediction of DNA N6- methyladenine site using deep learning-based approaches, Briefings in Bioinformatics 2022;23:bbac053.

21. Charoenkwan P, Nantasenamat C, Hasan MM et al. BERT4Bitter: a bidirectional encoder representations from transformers (BERT)-based model for improving the prediction of bitter peptides, Bioinformatics 2021:btab133.

22. Wu C, Gao R, Zhang Y et al. PTPD: predicting therapeutic peptides by deep learning and word2vec, BMC Bioinformatics 2019;20:456.

23. Zhang YF, Wang X, Kaushik AC et al. SPVec: A Word2vec-Inspired Feature Representation Method for Drug-Target Interaction Prediction, Front Chem 2019;7:895.

24. Yang X, Yang S, Li Q et al. Prediction of human-virus protein-protein interactions through a sequence embedding-based machine learning method, Comput Struct Biotechnol J 2020;18:153–161.

25. Tsukiyama S, Hasan MM, Fujii S et al. LSTM-PHV: prediction of human-virus protein–protein interactions by LSTM with word2vec, Briefings in Bioinformatics 2021;22:bbab228.

26. Yang X, Lian X, Fu C et al. HVIDB: a comprehensive database for human-virus protein-protein interactions, Brief Bioinform 2021;22:832–844.

27. The UniProt Consortium. UniProt: the universal protein knowledgebase, Nucleic Acids Res 2017;45:D158–D169.

28. Henikoff S, Henikoff JG. Amino acid substitution matrices from protein blocks, Proceedings of the National Academy of Sciences of the United States of America 1992;89:10915–10919.

29. Oughtred R, Rust J, Chang C et al. The BioGRID database: A comprehensive biomedical resource of curated protein, genetic, and chemical interactions, Protein science : a publication of the Protein Society 2021;30:187–200.

30. Fu L, Niu B, Zhu Z et al. CD-HIT: accelerated for clustering the next-generation sequencing data, Bioinformatics 2012;28:3150–3152.

31. Devlin J, Chang M-W, Lee K et al. BERT: Pre-training of Deep Bidirectional Transformers for Language Understanding. 2018, arXiv:1810.04805.

32. Vaswani A, Shazeer N, Parmar N, et al. Attention Is All You Need. 2017, arXiv:1706.03762.

33. Wu H, Gu X. Max-Pooling Dropout for Regularization of Convolutional Neural Networks. 2015, arXiv:1512.01400.

34. Pedregosa F, Varoquaux G, Gramfort A et al. Scikit-learn: Machine Learning in Python. 2012, arXiv:1201.0490.

35. van der Maaten L, Hinton G. Visualizing Data using t-SNE, Journal of Machine Learning Research 2008;9:2579–2605.

